# VRPTR prediction of individual language activation and uncertainty from resting state fMRI

**DOI:** 10.64898/2026.09.24.754231

**Authors:** Daniel A. Di Giovanni, D. Louis Collins

## Abstract

Resting-state connectivity can predict task-evoked fMRI activation, but correspondence with an individual task map may partly reflect a shared population pattern. We evaluated the Variational Resting-state-to-Task Prediction TransformeR (VRPTR), a three-dimensional encoder-decoder combining a compressed Transformer bottleneck, variational latent sampling, and multiscale skip connections. Models were trained in 360 healthy adults from the WU-Minn Human Connectome Project and evaluated in 40 held-out participants from the same dataset for the story-versus-math language contrast. VRPTR achieved mean voxel-map Pearson r=0.642 and Dice AUC=0.519, exceeding compact volumetric BrainSurfCNN-like and SWIFUN-like comparators by Δr=0.0376 and 0.0335, respectively. A template fixed from the 360 training maps yielded residual correlation 0.364; VRPTR improved correlation but not mean absolute or root mean squared error over this template. A matched factorial ablation associated Transformer processing with improved map correlation (Δr=0.0168); the variational main effect did not survive multiple-comparison correction. Component swaps indicated that subject-specific activation patterns depended more on multiscale skip features than on the bottleneck representation. Latent sampling produced a modest association between posterior standard deviation and voxelwise absolute error (mean Spearman ρ=0.091). Raw 95% intervals covered only 4.7% of observed values. Five-fold calibration within the held-out cohort increased coverage to 94.8%, and spatially varying uncertainty modestly improved probabilistic scores over a constant-width model. Uncertainty also increased under input degradation. These findings support the contribution of Transformer processing to language-map prediction and distinguish it from subject-specific spatial conditioning and relative uncertainty estimation. Independent validation of calibrated uncertainty and evaluation in clinical cohorts remain necessary.

## 1. Introduction

Task-evoked functional MRI maps distributed cognitive systems, but task acquisitions depend on successful performance and are vulnerable to motion, fatigue, neurological impairment, limited test-retest reliability, and disease-related changes in neurovascular coupling (D’Esposito et al., 2003; Elliott et al., 2020). Resting-state fMRI is easier to acquire, and intrinsic network architecture is broadly preserved across rest and task; activity propagated over resting-state networks can also predict task activation (Cole et al., 2014; Cole et al., 2016). This has motivated methods that infer task-evoked activation from task-free connectivity, beginning with multivariate prediction from resting-state fingerprints and extending to surface convolutional and Transformer architectures (Kwon et al., 2025; Ngo et al., 2022; Tavor et al., 2016). The translational motivation is especially clear for eloquent-cortex mapping, where task performance can be difficult and resting-state methods may provide complementary information (Di Giovanni & Collins, 2023).

Language activation is a demanding benchmark because it is distributed across multiple cortical regions, varies in lateralization and spatial expression across individuals, and depends on successful paradigm performance. Language fMRI also has direct scientific and clinical value because it supports assessment of hemispheric dominance, localization of eloquent cortex, characterization of functional reorganization; all important in presurgical planning in tumor and epilepsy surgery (Binder et al., 1996; Bizzi et al., 2008; Szaflarski et al., 2017). For these reasons, the story-versus-math contrast provides a stringent and clinically motivated target for individualized prediction and uncertainty, particularly when a patient cannot perform a conventional language task reliably (Barch et al., 2013; Benjamin et al., 2017; Di Giovanni & Collins, 2023; Kumar et al., 2020).

The scientific challenge is not simply to generate a plausible task map. Canonical task contrasts contain strong shared anatomy, so relatively high whole-map correspondence can be achieved by approximating a cohort-average activation pattern. Predictions can therefore regress toward the cohort mean while missing individual variation. A complementary test is whether a model recovers each participant’s deviation from that template. This distinction is central to individual-differences neuroimaging because group averaging can obscure reproducible person-specific topography and connectivity (Amico & Goñi, 2018; Gordon et al., 2017; Laumann et al., 2015); resting connectivity is highly person-specific (Finn et al., 2015; Mueller et al., 2013), and stable individual factors persist across rest and task states (Gratton et al., 2018). Rest-to-task models should therefore be evaluated both for global reconstruction and for individualized residual recovery.

The **V**ariational **R**esting-state-to-**T**ask **P**rediction **T**ransforme**R** (VRPTR) was originally introduced as a surface-based architecture for clinical functional MRI (Di Giovanni, 2024). The architecture combines convolutional encoding, a Transformer bottleneck (Vaswani et al., 2017), and a variational latent representation to address the nonlinear mapping from intrinsic connectivity to task activation. The present study builds on that concept as a controlled volumetric experiment using three-dimensional convolutions and asks which architectural components account for prediction, individualization, and probabilistic utility. This choice was deliberate since presurgical functional maps are interpreted in direct correspondence with anatomical MRI for localization, lateralization, and surgical planning, and a volumetric prediction can therefore be visualized and integrated without an additional surface-projection step. We therefore prioritized compatibility with image-based clinical workflows and whole-brain prediction over the geometric advantages offered by cortical surface models.

Three questions motivate the study. First, does a compressed Transformer bottleneck improve prediction of observed language-task activation? Second, how do bottleneck and multiscale convolutional features contribute to subject-specific activation patterns? Third, does repeated latent sampling provide useful information about prediction error? The main evaluation concerns correspondence between predicted and observed voxel maps. Secondary analyses examine deviations from a population template, participant matching, and component swaps to interpret that performance. We use point prediction to mean one predicted task map per participant, obtained by decoding the latent mean (z = μ); this differs from averaging maps produced by repeated latent draws. A matched Transformer-by-variational ablation, posterior sampling, cross-fitted calibration, and controlled input degradation address the architectural and uncertainty questions.

## 2. Methods

### 2.1. Participants and prediction task

Healthy-adult data were drawn from the WU-Minn Human Connectome Project, as described by Van Essen et al. (2013). A fixed cohort of 360 participants was used for model development and training, and 40 additional healthy HCP participants were reserved for evaluation. The 40 held-out participants were not used for network fitting, checkpoint selection, or latent ranking. Uncertainty calibration was subsequently assessed by five-fold cross-fitting within this cohort (Section 2.8), so its calibrated uncertainty results constitute internal validation. The task target was the HCP story-versus-math language contrast (Barch et al., 2013). We restricted the experiment to this single functional domain a priori so that architecture, individualization, and uncertainty could be interpreted against one coherent and clinically salient target rather than averaged across tasks with different networks, signal distributions, and behavioral demands. Language provides a stringent test because prediction must recover both a distributed group-level network and individual variation in its spatial expression and lateralization.

### 2.2. Resting-state fingerprints and task maps

All inputs and outputs were represented in volumetric image space. The volumetric formulation retains a common volumetric grid rather than restricting prediction a priori to the cortical sheet, with prediction and evaluation restricted to the analysis mask. Potential advantages for subcortical structures or distorted anatomy require separate evaluation.

Functional images were processed from the available HCP resting-state runs and language-task runs using a custom Nipype workflow built from FSL and Nilearn components (Abraham et al., 2014; Jenkinson et al., 2012). The participant’s T1-weighted image was brain-extracted with BET using a fractional intensity threshold of 0.30 and aligned to the 2-mm MNI152 brain template with 12-degree-of-freedom FLIRT registration, normalized mutual information, and trilinear interpolation (Jenkinson & Smith, 2001). For each fMRI run, slice-timing correction was applied with TR = 0.72 s and the third image axis specified as the slice direction, followed by rigid-body motion correction with MCFLIRT. A mean EPI image was aligned to the brain-extracted T1 image using six-degree-of-freedom FLIRT registration with normalized mutual information, 640 histogram bins, and trilinear interpolation. The EPI-to-T1 and T1-to-MNI transforms were concatenated and applied to the motion-corrected four-dimensional run (Jenkinson et al., 2002; Jenkinson & Smith, 2001). Images were then smoothed with a 6-mm full-width-at-half-maximum Gaussian kernel, temporally detrended, voxelwise standardized, and high-pass filtered at 0.01 Hz; no temporal low-pass filter was applied at this stage. The custom workflow did not include a separate susceptibility-distortion correction, global signal regression, or an additional nuisance-regressor model beyond rigid-body motion correction and temporal detrending/filtering.

Resting-state connectivity fingerprints were constructed from all available preprocessed resting-state runs. The Schaefer 100-parcel, 17-network cortical atlas was used at 2-mm resolution (Schaefer et al., 2018). Parcel-mean and MNI152 gray-matter voxel time series were extracted separately from each run with 0.01-0.10 Hz temporal filtering, concatenated across runs, and standardized across the combined time axis. For each parcel, Pearson correlation with every gray-matter voxel produced one voxelwise seed-connectivity map. Stacking the 100 seed maps yielded the 100-channel volumetric connectivity fingerprint. The resulting feature array can be written as 100 × V, where V is the number of gray-matter voxels, and stored as a four-dimensional image with 100 feature volumes. The atlas defines the seeds used to construct the inputs; predicted activation and Dice overlap are evaluated voxelwise.

Language-task events were loaded separately for story and math blocks from the HCP event files and paired with the corresponding fMRI runs. A design matrix was constructed for each run using the SPM hemodynamic response function, cosine drift regressors, and a 0.01-Hz high-pass filter. Runs were fitted jointly with Nilearn’s GLM using an autoregressive noise model, a slice-time reference of 0.5, and the MNI152 gray-matter mask (Abraham et al., 2014). Because spatial smoothing had already been applied during preprocessing, no additional first-level smoothing was used. The prediction target was the story-minus-math effect-size contrast, saved as a voxelwise beta map in the same 2-mm MNI152 space as the resting-state fingerprints.

Each participant was thus represented by a volumetric resting-state connectivity fingerprint with 100 feature maps. Each feature map encodes connectivity between each gray-matter voxel and one reference parcel, producing a spatially resolved description of large-scale functional organization. The feature dimension indexes reference parcels, not time points or successive connectivity matrices. The task target was the voxelwise story-versus-math beta map in the same analysis space. The analysis mask retained voxels with finite targets and finite, nonzero fingerprints. Each target map was standardized to zero mean and unit population standard deviation within its mask; voxels outside the mask were excluded from losses and metrics. All models received identical inputs and targets.

### 2.3. VRPTR architecture

VRPTR is a four-level three-dimensional residual encoder-decoder with a compulsory transformed bottleneck (Figure 1), drawing on residual learning and U-Net-like encoder-decoder designs (He et al., 2016; Ronneberger et al., 2015). The encoder uses base width 32 and doubles channels with spatial downsampling. Shallower encoder features are retained as multiscale skips, but the deepest encoder tensor is excluded from the skip set; it has no direct route to the decoder. At the deepest resolution, the encoder produces 256 feature maps. Each feature map is flattened and compressed to a 32-element vector, yielding 256 compact feature tokens. These tokens are processed jointly by a six-layer Transformer encoder with eight attention heads, allowing information from different resting-state connectivity features to be integrated before task-map reconstruction. Rather than assigning each token a single fixed latent representation, the variational component describes each latent value as a probability distribution. Two linear layers parameterize the mean, *μ_φ_*(*x*), and standard deviation, *σ_φ_*(*x*), of that distribution from the encoded input *x*:

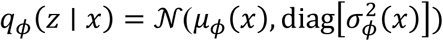

**Figure 1.**
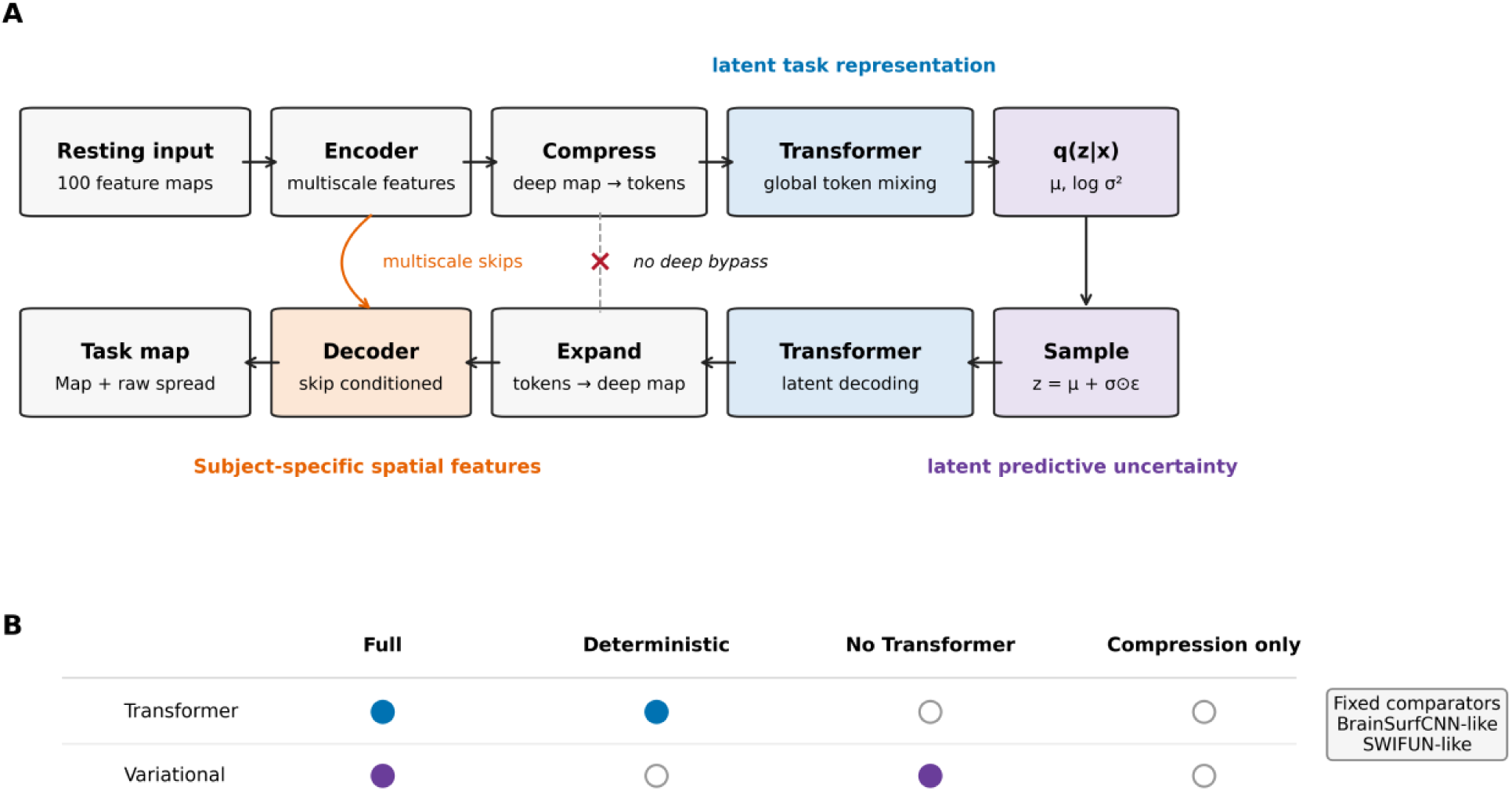
VRPTR architecture and factorial study design. (A) A resting-state fingerprint containing 100 seed-connectivity feature maps enters the three-dimensional encoder. The deepest representation is compressed into feature tokens, processed by a Transformer encoder, parameterized as a latent distribution, sampled, processed by a Transformer decoder, expanded, and reconstructed with shallower multiscale skips. The deepest encoder tensor has no direct skip connection to the decoder. A point prediction decodes z=μ; repeated latent draws provide raw predictive spread. (B) The four variants cross Transformer processing with variational sampling. BrainSurfCNN-like and SWIFUN-like are compact volumetric comparators.

Here, *q_φ_*(*z* ∣ *x*) is the model’s approximate posterior distribution over plausible latent representations *z* for the observed resting-state input. The mean specifies the most likely latent representation, while the standard deviation describes how much that representation can vary. The covariance is diagonal, meaning that each latent element is assigned its own variance without explicitly modeling covariance between latent elements. A latent representation is then sampled using the reparameterization rule (Kingma & Welling, 2019):

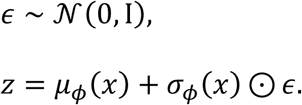

In practical terms, the model first draws random noise ***ε*** from a standard normal distribution. This noise is scaled by the predicted standard deviation and added to the predicted mean. Repeating this process for the same resting-state input produces slightly different but plausible latent representations, which can be decoded into a distribution of predicted task maps rather than a single deterministic output.

The sampled latent tokens are subsequently processed by a six-layer Transformer decoder, expanded back to the original bottleneck dimensions, and passed through a three-level convolutional decoder. At each decoding level, the model incorporates the corresponding shallower encoder features through multiscale skip connections. A final **1** × **1** × **1** convolution converts the reconstructed features into the predicted task beta map. This volumetric formulation preserves the central architectural logic of the original surface-based VRPTR model while enabling prediction and evaluation directly in voxel space (Di Giovanni, 2024).

### 2.4. Comparators and factorial ablations

BrainSurfCNN-like and SWIFUN-like are compact volumetric adaptations of surface-convolutional and shifted-window Transformer approaches (Liu et al., 2021) to rest-to-task prediction (Kwon et al., 2025; Ngo et al., 2022). They are controlled adaptations rather than exact reproductions. All learned models used the same volumetric fingerprints, targets, split, and evaluation pipeline.

A matched 2 × 2 factorial design crossed Transformer processing with variational sampling. Full VRPTR contained both components. Deterministic VRPTR retained the Transformer encoder-decoder but used *z = μ* and omitted the variance/KL pathway. No-Transformer VRPTR retained spatial compression, latent sampling, and expansion but replaced both Transformer stages with identity mappings. Compression-only VRPTR removed both Transformer processing and variational sampling. The four variants were trained from scratch with matched data, architecture scale, optimization, duration, and seed.

### 2.5. Training

VRPTR variants were trained for 100 epochs using AdamW (learning rate 2 × 10⁻⁵; weight decay 10⁻⁴), global batch size two, dropout 0.15, gradient clipping at 1.0, mixed precision, and random seed 42. The reconstruction objective was masked smooth L1 loss. From epoch 51, a subject-contrastive term (weight 0.08; temperature 0.20) encouraged each prediction to match its own target more strongly than targets from other participants in the distributed batch. Variational variants used a KL weight of 10⁻⁴, a 20-epoch deterministic warm-up, a 10-epoch latent-noise ramp, and a 30-epoch KL ramp. The prespecified final-epoch checkpoint was used for evaluation. Using the same fixed training duration and final checkpoint gave the four ablations a common optimization schedule without selecting epochs on the test cohort. This design does not establish that 100 epochs was optimal or that all variants had converged.

### 2.6. Task-map prediction and secondary analyses of subject-specific activation

The main prediction outcome was voxel-map Pearson correlation between each predicted and observed task map. Mean absolute error (MAE) and root mean squared error (RMSE), in standardized task-map units, additionally assessed absolute agreement. Dice AUC summarized overlap among the top 5%–50% of voxels ranked by absolute map magnitude, in 5-percentage-point steps, using trapezoidal integration divided by 0.45. Thus, Dice included both positive and negative contrast values rather than positive activation alone. Metrics used paired finite voxels; an empty mask yielded an undefined Dice score, and each nonempty mask retained at least one voxel at every threshold.

Secondary analyses assessed performance relative to a population template and recovery of participant-specific deviations. A single template T_train was the voxelwise mean of the 360 training participants’ standardized task maps, without restandardizing the mean. This template was fixed before test evaluation and used for all models. Template gain was r(prediction, observed map) − r(T_train, observed map). Observed and predicted residuals were the corresponding maps minus the same T_train (Figure 2A). No test task map contributed to this reference. A separate sensitivity analysis used the leave-one-out mean of the other 39 test maps; the prediction networks were not refitted.

**Figure 2.**
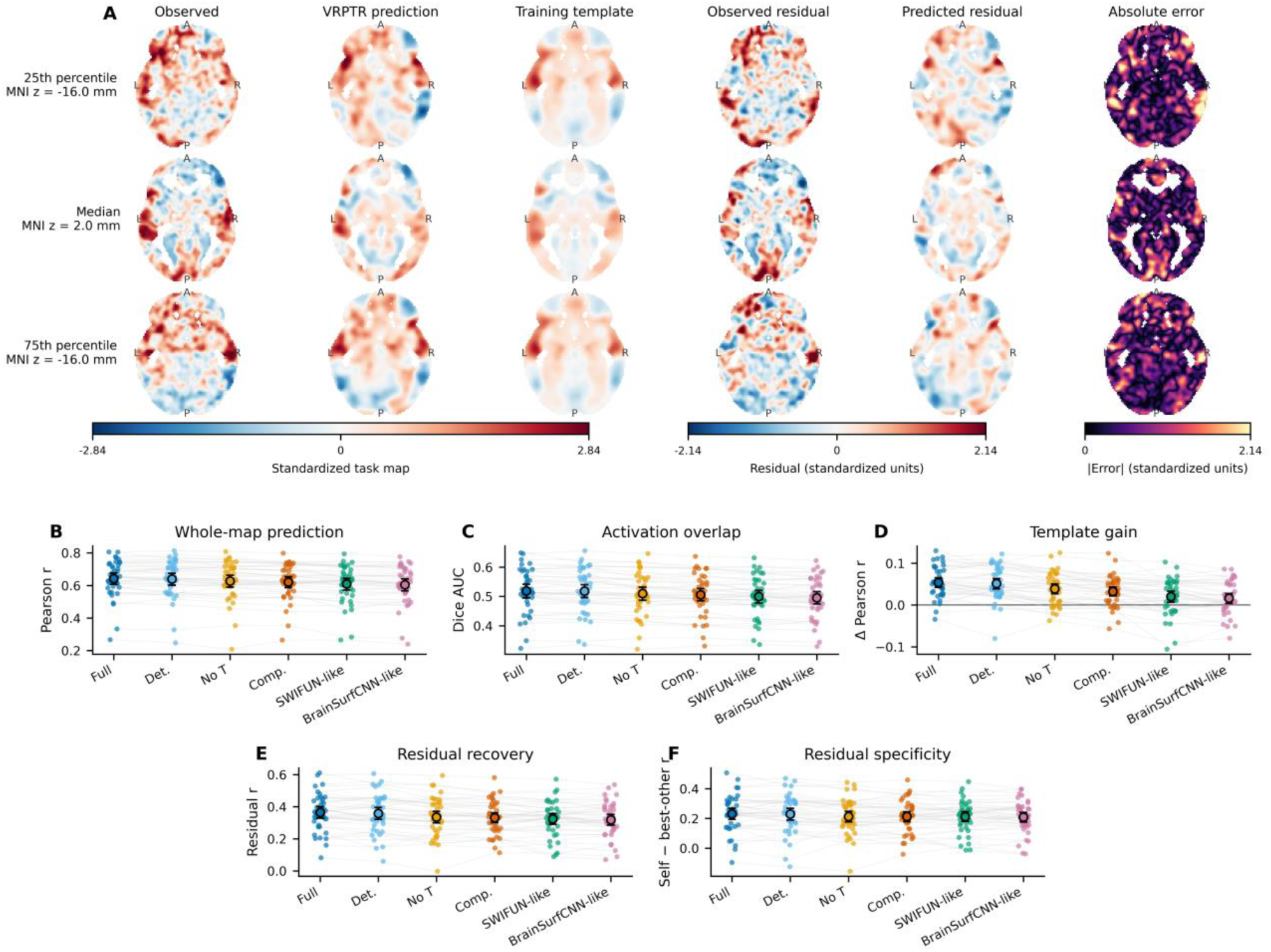
Observed and predicted language-task maps and secondary residual recovery. (A) Observed maps, full-VRPTR point predictions (z=μ), the fixed training-derived template, observed and predicted residuals, and absolute error. Rows show participants nearest the 25th, 50th, and 75th percentiles of whole-map Pearson correlation, selected before visual inspection: r=0.595, 0.650, 0.723, respectively. Slice coordinates and affine-derived orientation appear in the panels. Maps share a mask and fixed numeric scales within map classes; units are within-participant standardized task-map values. (B–F) Whole-map Pearson correlation, Dice AUC, correlation gain over the training template, residual correlation, and residual self-versus-best-other margin. Gray lines connect the same participant; black markers and bars show means and percentile bootstrap 95% confidence intervals (5,000 resamples). Residuals subtract the same training template from observed and predicted maps. Full, full VRPTR; Det., deterministic; No T, no-Transformer; Comp., compression-only. Panel B is primary; panels C–F are complementary or secondary.

**Figure 3.**
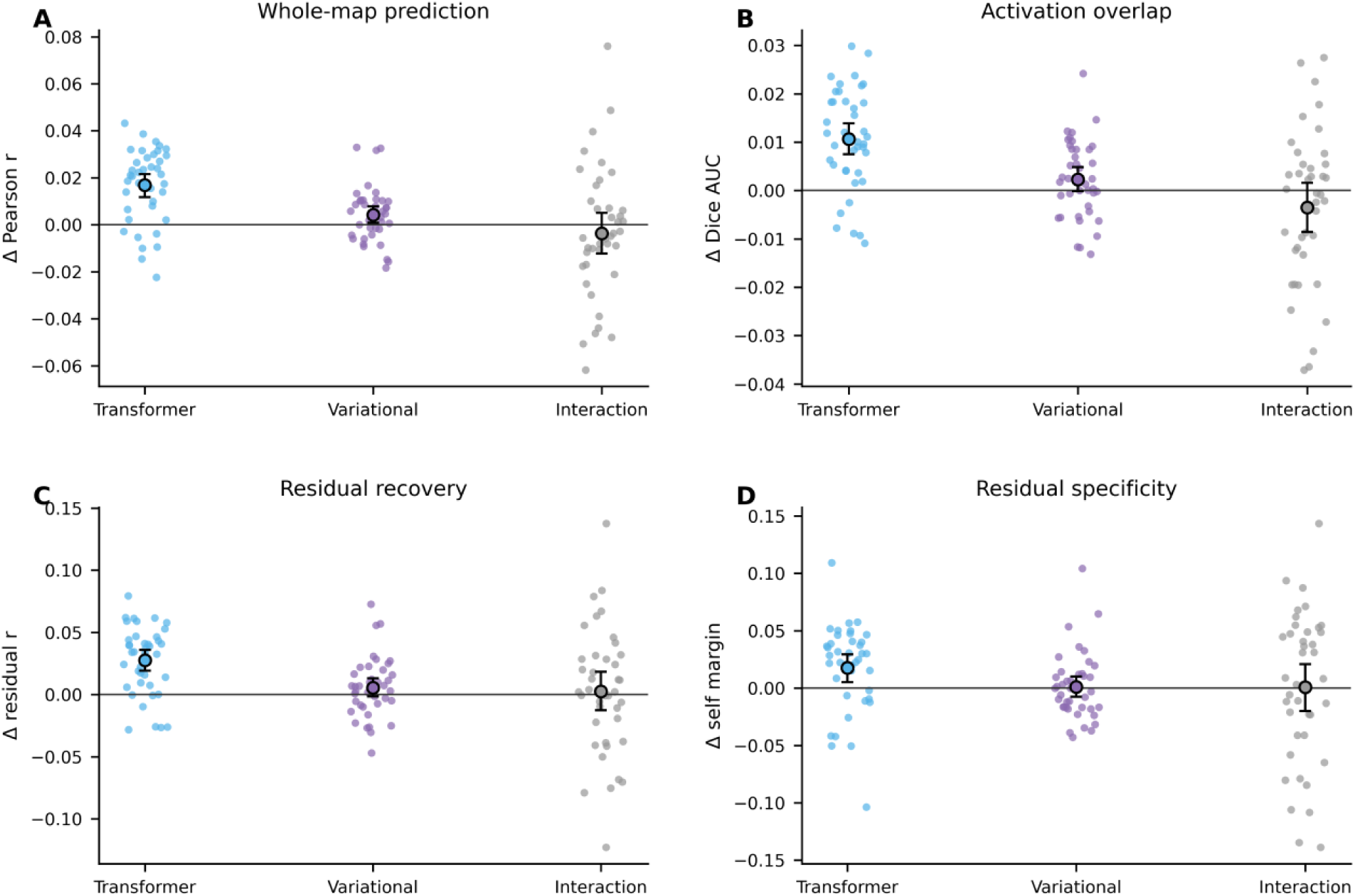
Factorial effects of Transformer processing and variational sampling. Panels A–D show whole-map Pearson correlation, Dice AUC, residual correlation, and residual self-versus-best-other margin. Points represent held-out participants; black markers and bars show means and percentile bootstrap 95% confidence intervals (5,000 resamples). Intervals are unadjusted; multiple-comparison-adjusted inference is reported in Table 2.

Specificity of participant-level predictions was further assessed by correlating each predicted residual map with every participant’s observed residual map. Residual self-versus-best-other margin was the correlation with the correct participant minus the largest correlation with any other participant. The same fixed training template was subtracted for every self and other comparison, with matching computed over the common finite-voxel mask. Top-1 matching required the correct participant to be the unique best match; ties were not counted as correct. A positive self-versus-best-other margin indicates a closer match to the correct participant than to any alternative. Because participant matching approached ceiling, residual correlation and this margin provide more informative secondary summaries. These analyses assess specificity within this cohort; they do not assume that every individual has a unique functional map or replace prediction accuracy as the main outcome.

To make spatial performance directly interpretable, Figure 2A shows full-VRPTR single voxel-map predictions obtained by decoding the latent mean (z = μ) for the held-out participants nearest the 25th, 50th, and 75th percentiles of full-VRPTR Pearson correlation. For each participant, the observed task map, point prediction, fixed training-derived population template, observed and predicted individualized residuals, and absolute error are displayed at the same axial slice.

### 2.7. Factorial and component-swap analyses

The four factorial conditions were full VRPTR, deterministic VRPTR, no-Transformer VRPTR, and compression-only VRPTR. For each participant and metric, the Transformer effect was one half of [(full − no-Transformer) + (deterministic − compression-only)]. The variational effect was one half of [(full − deterministic) + (no-Transformer − compression-only)]. The interaction was (full − deterministic) − (no-Transformer − compression-only). Whole-map correlation and Dice AUC summarized prediction performance; residual correlation and self-versus-best-other margin were secondary outcomes.

Component swaps tested which internal features supported subject-specific activation patterns. For each ordered pair of held-out participants, the recipient was the participant whose original prediction provided the reference, and the donor was another participant supplying replacement features. A bottleneck swap decoded the donor’s bottleneck representation together with the recipient’s multiscale skip features. A skip swap decoded the recipient’s bottleneck together with the donor’s skip features. These hybrids were compared with the recipient’s original prediction and with the donor’s and recipient’s observed task maps, using both whole maps and residual maps. All 40 × 39 = 1,560 ordered pairs were evaluated for full VRPTR. Effects were averaged across the 39 donors for each recipient before group inference, leaving 40 participant-level summaries rather than treating the pairs as independent observations. Donor shift was the hybrid’s donor-minus-recipient correlation minus the same difference for the recipient’s unmodified prediction. Residual shifts used the fixed training task-map template throughout. Additional interventions replaced skip features with their mean across the 40 test participants or decoded the mean latent representation from those participants to replace the bottleneck. These test-cohort feature interventions were descriptive and distinct from the training-derived task-map reference.

### 2.8. Posterior predictive quality and utility

For the full model, repeated latent draws induce a predictive distribution over task maps conditional on fixed network weights and fixed subject-specific skip features. We therefore refer to a latent posterior predictive distribution rather than a full Bayesian posterior. For each held-out participant, 128 latent samples were decoded to estimate the posterior mean, voxelwise posterior standard deviation, and central prediction intervals of the predicted task maps. Uncertainty utility was evaluated by the within-participant Spearman association between posterior standard deviation and absolute voxelwise error, by the reduction in error among retained voxels after excluding the highest-uncertainty output voxels from evaluation, and by Gaussian negative log-likelihood and continuous ranked probability score (CRPS) (Gneiting & Raftery, 2007). Latent sampling represents variation conditional on fixed weights and skip features. It does not explicitly account for all sources of predictive error, including measurement noise in the observed task map or uncertainty in the fitted weights and deterministic pathways. We therefore calibrated the raw posterior standard deviation before interpreting it as predictive uncertainty. Uncertainty metrics used a reproducible subset of up to 25,000 valid output voxels per participant. Selective prediction ranked these voxels by posterior standard deviation and retained 25%, 50%, 75%, 90%, and 100%. Integrated gain was one minus the normalized trapezoidal area under this MAE–coverage curve divided by MAE over all sampled voxels. It summarizes the full tested coverage range, not a single withholding threshold. Only the scored output voxels changed; inputs and predictions were unchanged. For each participant and voxel, the calibrated predictive variance was defined as follows (Levi et al., 2022; Mehrtash et al., 2020):

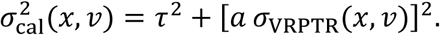

Here, *x* denotes the participant’s resting-state input and *v* denotes a voxel. The quantity *σ*_VRPTR_(*x*, *v*) is the raw standard deviation of the task-map predictions obtained by repeatedly sampling the VRPTR latent representation. The parameter *a* rescales this raw uncertainty so that differences between relatively certain and uncertain voxels are expressed on the scale of the observed prediction error. The parameter *τ* is an additional uncertainty floor shared across voxels. It represents prediction error that is not explained by variation in the latent samples.

The model therefore combines two forms of uncertainty. The term *τ*^2^ accounts for a baseline level of error that is present throughout the predicted map, whereas [*a σ*_VRPTR_(*x*, *v*)]^2^ allows uncertainty to vary spatially according to the model’s latent posterior. Voxels with larger raw VRPTR variability receive wider calibrated prediction intervals, but even voxels with very little latent variability retain the baseline uncertainty represented by *τ*.

Calibration was performed using five-fold cross-fitting across the 40 held-out participants. In each fold, the scaling parameters *a* and *τ* were estimated from 32 participants and then applied to the remaining eight. This procedure was repeated until every participant had been evaluated using calibration parameters estimated entirely from other participants. It therefore prevented the observed task map of a participant from being used to calibrate that participant’s own uncertainty. The two positive parameters minimized Gaussian negative log-likelihood over up to 500,000 pooled calibration voxels; the constant-width baseline was fitted on the same folds. Network weights remained fixed during this step. Although each participant was evaluated out of fold, the calibrator learned from other members of the same held-out cohort; these results do not establish calibration on an independent external dataset. Deployment would require fitting the calibration parameters on a separate calibration set and freezing them before final evaluation.

To test whether the spatial pattern of VRPTR uncertainty added useful information, we compared this calibrated heteroscedastic model with a simpler homoscedastic model. The heteroscedastic model used both *a* and *τ*, allowing prediction intervals to vary from voxel to voxel. The homoscedastic model used only *τ*, assigning the same uncertainty width to every voxel. Better probabilistic scores for the heteroscedastic model would therefore indicate that the spatial variation in VRPTR posterior uncertainty was useful beyond a single global error estimate. We additionally used 48 latent draws and up to 10,000 valid voxels per participant under Gaussian noise (standard deviations 0.25, 0.50, 1.00 in standardized input units) and whole input feature-map dropout (10%, 25%, 50%). Perturbations were nested within participant and corruption family to test whether uncertainty increased as inputs degraded. This analysis evaluated uncertainty awareness, not improved robustness of the posterior mean.

### 2.9. Statistical inference

All model comparisons were paired across the same 40 participants. Mean effects were summarized with participant-level bootstrap 95% confidence intervals using 20,000 resamples and two-sided sign-flip permutation tests using 20,000 permutations (Winkler et al., 2014). Benjamini-Hochberg false-discovery-rate correction was applied within prespecified analysis families (Benjamini & Hochberg, 1995). Voxels were not treated as independent group-level observations; voxelwise quantities were summarized within participant before inference. These intervals and tests describe variation across the evaluated participants conditional on the fitted networks. A single training seed does not quantify variability across independent training runs, and overlapping calibration folds and shared swap donors can induce dependence between participant-level summaries. MAE and RMSE were descriptive additions and were not added to the existing multiple-testing families. Figure confidence intervals used 5,000 bootstrap resamples.

## 3. Results

### 3.1. VRPTR improved held-out task-map prediction

VRPTR achieved the highest mean correlation and Dice AUC among the evaluated models in the 40 held-out healthy HCP participants (Pearson r=0.642; Dice AUC=0.519; Table 1). Relative to BrainSurfCNN-like, it increased correlation by 0.0376 (95% CI 0.0330–0.0422; p=5 × 10⁻⁵) and was better in all 40 participants. Relative to SWIFUN-like, it increased correlation by 0.0335 (95% CI 0.0266–0.0406; p=5 × 10⁻⁵) and was better in 39 of 40 participants. Activation-overlap improvements were concordant (Figure 2B–C). Figure 2A displays algorithmically selected lower-quartile, median, and upper-quartile examples, including the observed and predicted task maps, the population template, individualized residuals, and absolute error.

**Table 1.**
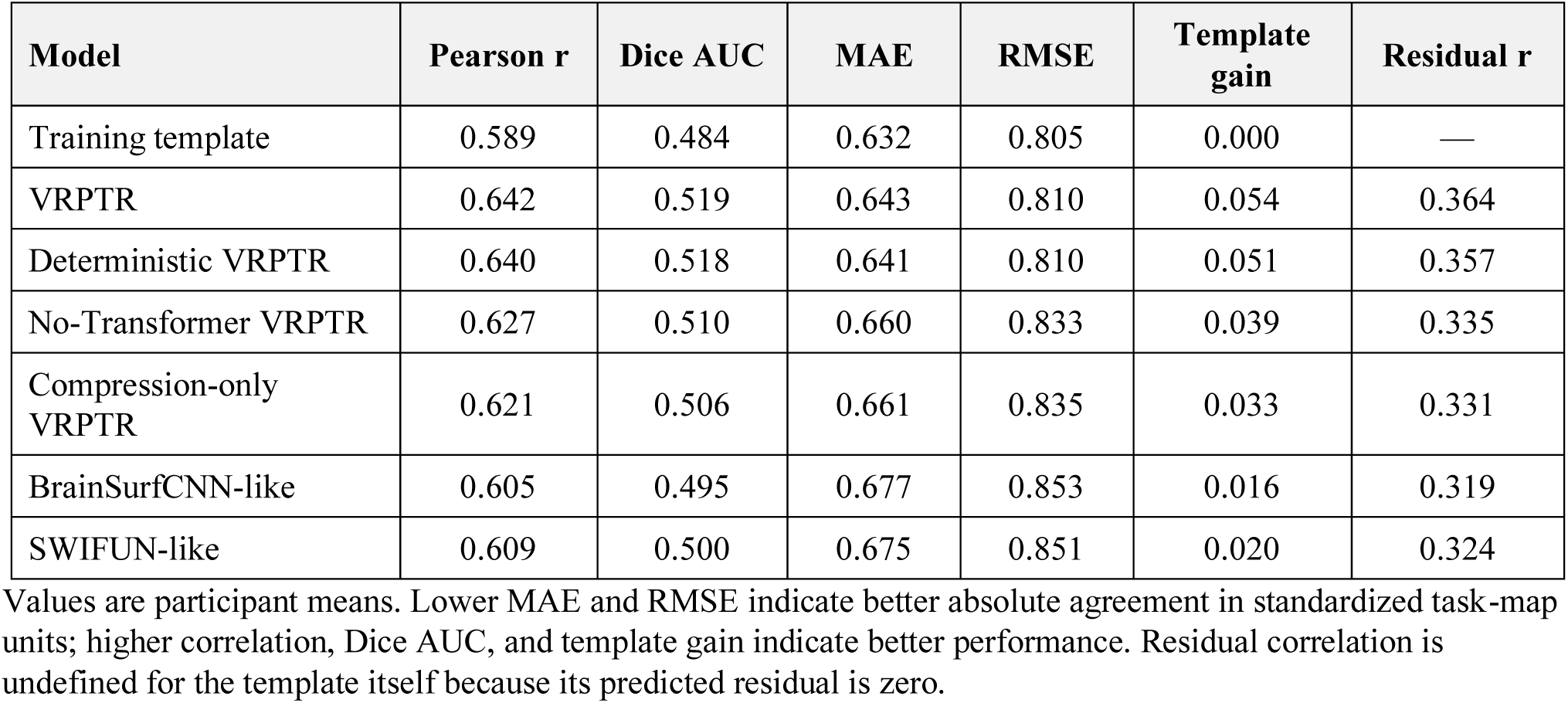
Prediction and secondary residual analyses in 40 held-out healthy HCP participants.

| Model | Pearson r | Dice AUC | MAE | RMSE | Template gain | Residual r |
| --- | --- | --- | --- | --- | --- | --- |
| Training template | 0.589 | 0.484 | 0.632 | 0.805 | 0.000 | — |
| VRPTR | 0.642 | 0.519 | 0.643 | 0.810 | 0.054 | 0.364 |
| Deterministic VRPTR | 0.640 | 0.518 | 0.641 | 0.810 | 0.051 | 0.357 |
| No-Transformer VRPTR | 0.627 | 0.510 | 0.660 | 0.833 | 0.039 | 0.335 |
| Compression-only VRPTR | 0.621 | 0.506 | 0.661 | 0.835 | 0.033 | 0.331 |
| BrainSurfCNN-like | 0.605 | 0.495 | 0.677 | 0.853 | 0.016 | 0.319 |
| SWIFUN-like | 0.609 | 0.500 | 0.675 | 0.851 | 0.020 | 0.324 |

Values are participant means. Lower MAE and RMSE indicate better absolute agreement in standardized task-map units; higher correlation, Dice AUC, and template gain indicate better performance. Residual correlation is undefined for the template itself because its predicted residual is zero.

The training template achieved mean Pearson r=0.589 and Dice AUC=0.484. VRPTR improved these outcomes but did not reduce average absolute error: MAE was 0.643 versus 0.632 for the template, and RMSE was 0.810 versus 0.805. VRPTR had lower MAE in 16/40 participants and lower RMSE in 17/40. These descriptive results distinguish spatial-pattern correspondence from absolute agreement.

### 3.2. Secondary analyses supported recovery of subject-specific task structure

VRPTR predictions exceeded the training-derived template by a mean correlation of 0.0536, compared with 0.0160 for BrainSurfCNN-like and 0.0202 for SWIFUN-like. VRPTR exceeded the template’s correlation in 38/40 participants. Its mean residual correlation was 0.364, its residual self-versus-best-other margin was 0.231, and the correct observed residual map was the unique best match for 38/40 predictions. Compared with BrainSurfCNN-like, VRPTR increased residual correlation by 0.0452 (95% CI 0.0365–0.0536; q<0.001) and residual self-margin by 0.0221 (95% CI 0.0090–0.0344; q=0.004). Corresponding improvements over SWIFUN-like were 0.0401 (95% CI 0.0276–0.0528; q<0.001) and 0.0192 (95% CI 0.0036–0.0351; q=0.036). With a common template, between-model differences in template gain equal their whole-map correlation differences.

The test-cohort leave-one-out sensitivity analysis preserved the model ranking. For full VRPTR, template gain was 0.0652, residual correlation was 0.386, and residual top-1 matching was 39/40. The corresponding training-template estimates were smaller, but the conclusion that predictions recovered participant-specific residual structure was retained.

### 3.3. Transformer processing accounted for the architectural advantage

The matched factorial analysis supported a Transformer contribution within the matched architecture (Table 2). Averaged across variational conditions, Transformer processing alone increased correlation by 0.0168, Dice AUC by 0.0106, residual correlation by 0.0275, and residual self-margin by 0.0177. Each effect survived false-discovery-rate correction. The variational main effect on correlation was smaller (0.0043) and did not survive correction; variational effects on activation overlap and individualization were also nonsignificant. The Transformer-by-variational interaction was not significant.

**Table 2.**
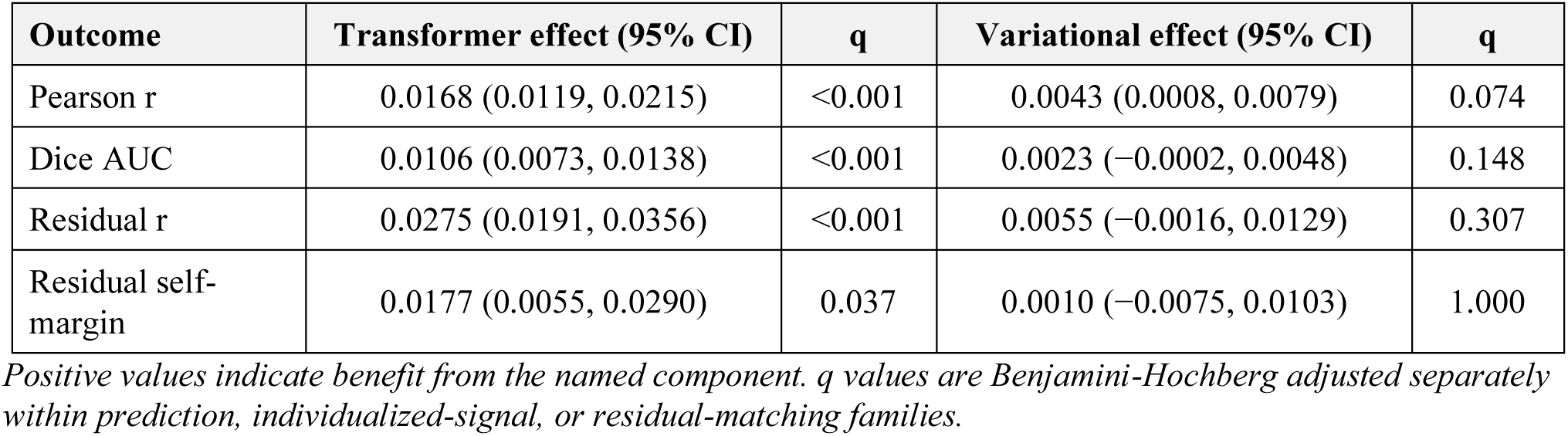
Matched factorial effects.

| Outcome | Transformer effect (95% CI) | q | Variational effect (95% CI) | q |
| --- | --- | --- | --- | --- |
| Pearson r | 0.0168 (0.0119, 0.0215) | <0.001 | 0.0043 (0.0008, 0.0079) | 0.074 |
| Dice AUC | 0.0106 (0.0073, 0.0138) | <0.001 | 0.0023 (−0.0002, 0.0048) | 0.148 |
| Residual r | 0.0275 (0.0191, 0.0356) | <0.001 | 0.0055 (−0.0016, 0.0129) | 0.307 |
| Residual self-margin | 0.0177 (0.0055, 0.0290) | 0.037 | 0.0010 (−0.0075, 0.0103) | 1.000 |

*Positive values indicate benefit from the named component. q values are Benjamini-Hochberg adjusted separately within prediction, individualized-signal, or residual-matching families.*

**Table 3.**
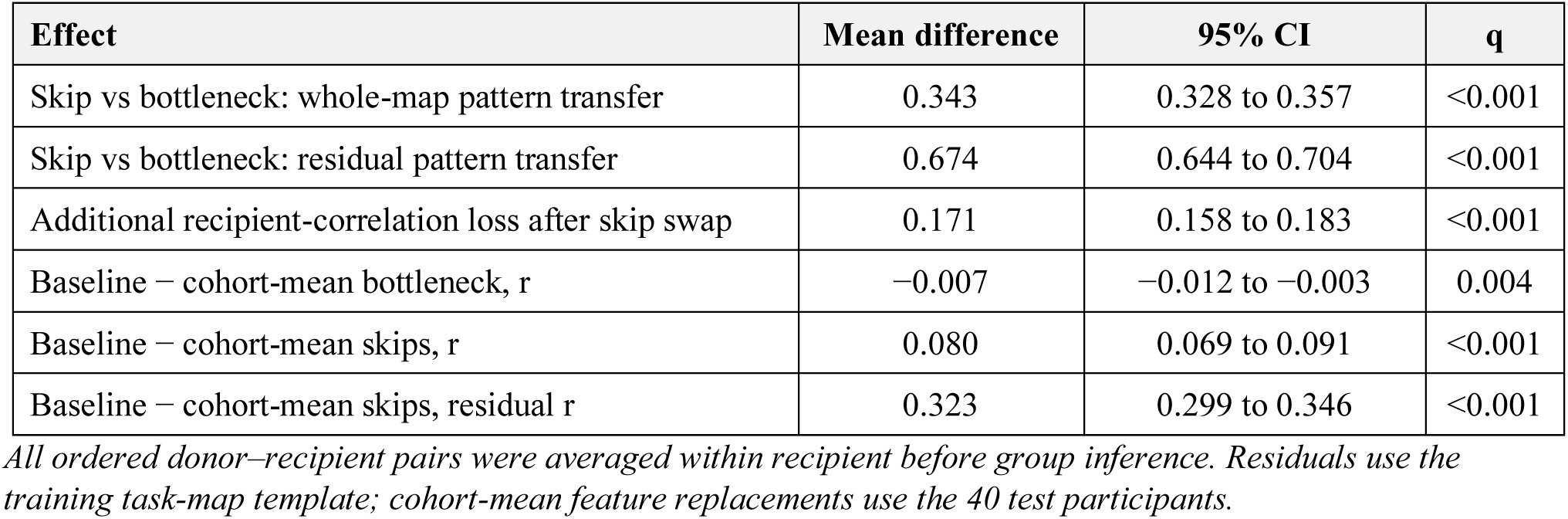
Component-swap effects in full VRPTR.

| Effect | Mean difference | 95% CI | q |
| --- | --- | --- | --- |
| Skip vs bottleneck: whole-map pattern transfer | 0.343 | 0.328 to 0.357 | <0.001 |
| Skip vs bottleneck: residual pattern transfer | 0.674 | 0.644 to 0.704 | <0.001 |
| Additional recipient-correlation loss after skip swap | 0.171 | 0.158 to 0.183 | <0.001 |
| Baseline – cohort-mean bottleneck, r | −0.007 | −0.012 to −0.003 | 0.004 |
| Baseline – cohort-mean skips, r | 0.080 | 0.069 to 0.091 | <0.001 |
| Baseline – cohort-mean skips, residual r | 0.323 | 0.299 to 0.346 | <0.001 |

*All ordered donor–recipient pairs were averaged within recipient before group inference. Residuals use the training task-map template; cohort-mean feature replacements use the 40 test participants.*

**Table 4.** Posterior predictive quality and utility.

| Analysis | Estimate | 95% CI or comparison | q |
| --- | --- | --- | --- |
| Uncertainty–absolute error Spearman $\rho$ | 0.091 | 0.078 to 0.105 | <0.001 |
| Integrated selective-prediction gain | 5.2% | 4.5% to 5.9% | <0.001 |
| Heteroscedastic NLL benefit | 0.0061 | 0.0037 to 0.0084 | <0.001 |
| Heteroscedastic CRPS benefit | 0.00083 | 0.00055 to 0.00113 | <0.001 |
| Raw nominal 95% coverage | 4.7% | Target 95% | — |
| Calibrated nominal 95% coverage | 94.8% | Target 95% | — |
| Noise uncertainty/degradation $\rho$ | 0.845 | Across severity | <0.001 |
| Feature-map-dropout uncertainty/degradation $\rho$ | 0.850 | Across severity | <0.001 |

*NLL and CRPS compare the cross-fitted calibrated heteroscedastic model with a cross-fitted constant-width model.*

Directly comparing the two Transformer models, full and deterministic VRPTR did not differ significantly in whole-map correlation (Δ=0.0024, 95% CI −0.0034 to 0.0082; p=0.43) or residual correlation (Δ=0.0068, 95% CI −0.0043 to 0.0180; q=0.330). The confidence interval includes effects in either direction; this comparison does not establish equivalence. The observed prediction advantage was retained by the deterministic Transformer model.

### 3.4. Multiscale skip features contributed most to subject-specific activation patterns

Component swaps changed predictions more strongly when the replacement features came through the skip pathways (Figure 4). Replacing recipient skips with those of another participant shifted predictions toward that donor’s activation pattern more than replacing the bottleneck did. The skip-minus-bottleneck difference was 0.343 for the whole-map transfer measure and 0.674 for the residual transfer measure (both q<0.001). Skip swaps also caused an additional 0.171 loss of correlation with the recipient’s observed map and a larger change from the original prediction (both q<0.001).

**Figure 4.**
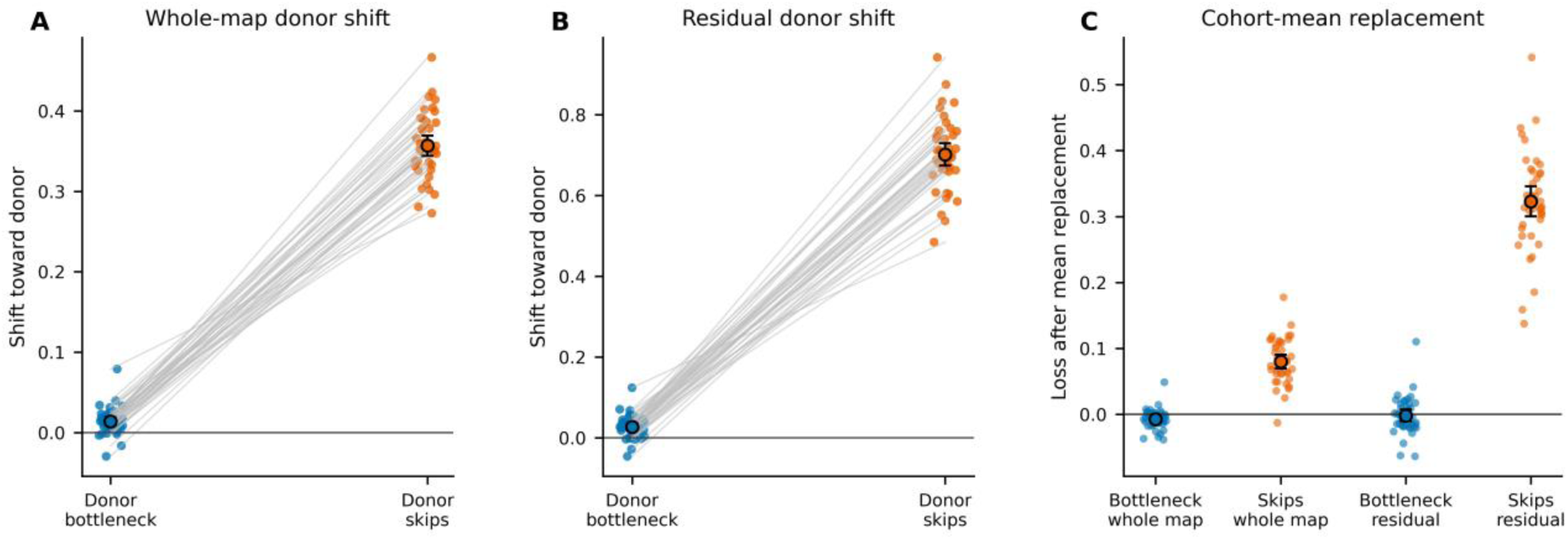
Effects of exchanging bottleneck and skip features in full VRPTR. The recipient supplies the reference prediction and the donor supplies replacement features (Section 2.7). (A–B) Donor shift is the change in donor-minus-recipient correlation relative to the recipient’s unmodified prediction, using whole maps or training-template residuals. (C) Correlation loss after replacing the bottleneck or skips with test-cohort mean features; these are descriptive interventions. Pairwise swaps were averaged over 39 donors within each of 40 recipients before inference. Black markers and bars show recipient means and percentile bootstrap 95% confidence intervals (5,000 resamples).

Replacing the subject-specific bottleneck with the test-cohort mean latent representation decoded through the bottleneck slightly improved mean correlation, whereas replacing subject-specific skips with cohort-mean skips reduced correlation by 0.080 and residual correlation by 0.323. These interventions are consistent with a relatively shared bottleneck representation and a stronger contribution of skip features to subject-specific spatial patterns.

### 3.5. Latent variability provided a modest relative uncertainty signal

The uncertainty analysis evaluated the spatial variability of sampled predictions (Figure 5). Within participants, posterior standard deviation had a modest association with voxelwise absolute error with mean Spearman ρ=0.091 (95% CI 0.078–0.105; q<0.001), and the association was positive in all 40 participants. The integrated selective-prediction gain across retained fractions of 25%–100% was 5.2% (95% CI 4.5%–5.9%; q<0.001), again with a positive effect in every participant.

**Figure 5.**
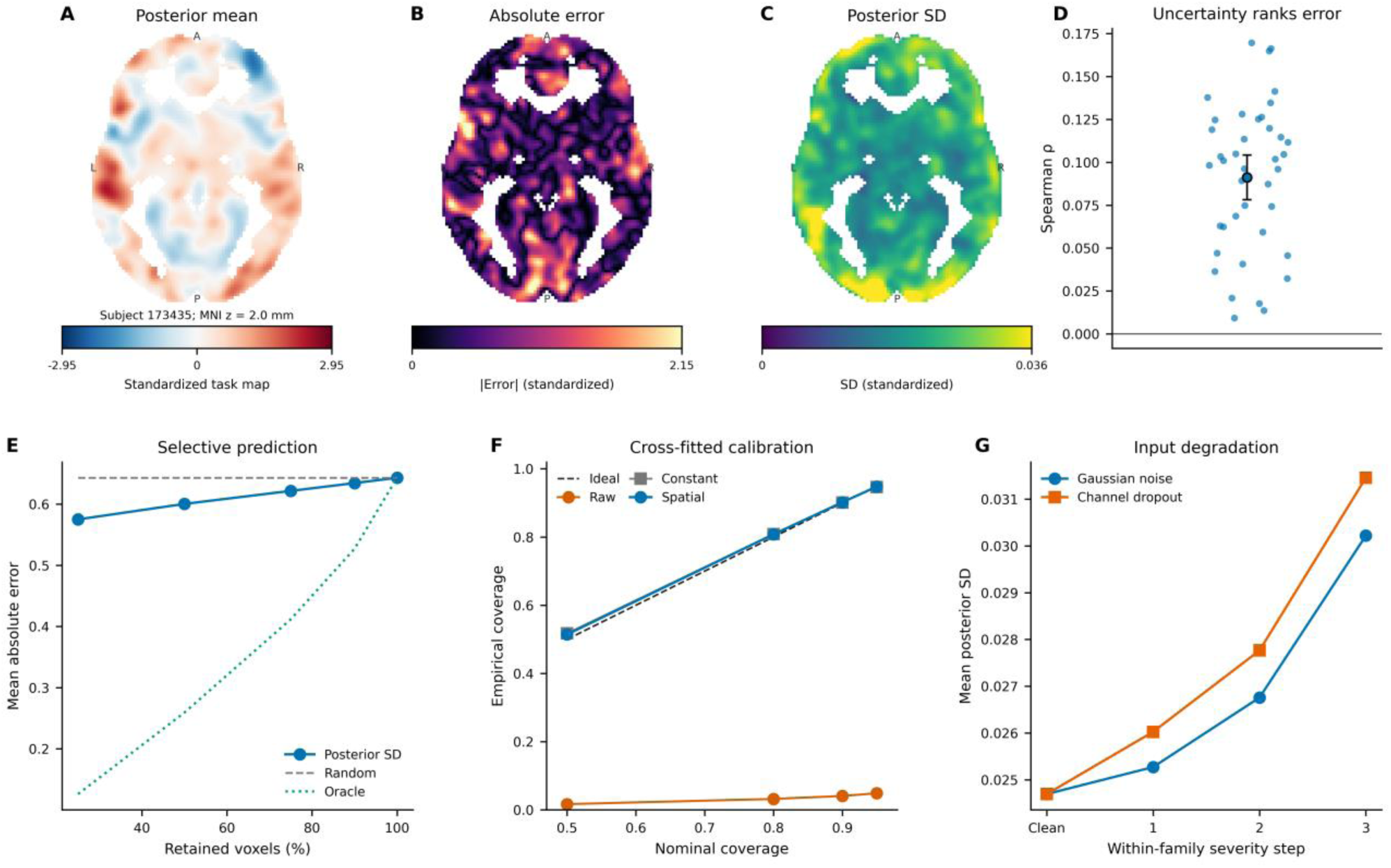
Latent variability and calibrated predictive uncertainty. (A–C) Posterior-mean map, absolute error, and standard deviation across 128 latent draws for the independently selected median-performance participant (173435; MNI z=2 mm), with numeric standardized-unit scales and affine-derived orientation. (D) Within-participant Spearman associations between posterior standard deviation and absolute error. (E) MAE among retained sampled output voxels, comparing uncertainty ranking, random selection, and observed-error ranking (oracle). The 5.2% summary integrates the curve over retained fractions of 25%, 50%, 75%, 90%, and 100%; inputs are unchanged. (F) Nominal versus empirical coverage for raw Gaussian intervals and five-fold cross-fitted constant-width and spatially varying calibration; raw Gaussian 95% coverage is 4.9%, compared with 4.7% for empirical latent-sample intervals reported in the text. (G) Mean posterior standard deviation by severity step: Gaussian noise SD 0, 0.25, 0.50, 1.00; input feature-map dropout 0%, 10%, 25%, 50%. These are different perturbation scales. Calibration was internal to the 40-participant evaluation cohort.

Raw intervals were markedly underdispersed: empirical coverage was 1.7%, 3.2%, 4.0%, and 4.7% for nominal 50%, 80%, 90%, and 95% intervals. Cross-fitted calibration required a mean multiplicative scale of 17.9 and a residual uncertainty floor of 0.676. After calibration, aggregate coverage was 80.8%, 90.2%, and 94.8% for nominal 80%, 90%, and 95% intervals; constant-width calibration also reached 94.7% coverage at the 95% level. Spatially calibrated 95% coverage ranged from 82.9% to 98.8% across participants. The calibrated heteroscedastic model improved negative log-likelihood over a calibrated constant-width model by 0.0061 (95% CI 0.0037–0.0084; q<0.001) and CRPS by 0.00083 (95% CI 0.00055–0.00113; q<0.001). Spatial posterior variation therefore carried information beyond a single global error bar, but raw posterior spread could not be interpreted as a calibrated prediction interval.

### 3.6. Posterior uncertainty tracked input degradation

Posterior uncertainty increased monotonically under both corruption families. Across severity levels within participants, uncertainty tracked prediction degradation with mean Spearman ρ=0.845 for Gaussian noise and ρ=0.850 for input feature-map dropout (both q<0.001). At the highest Gaussian-noise severity, mean posterior standard deviation rose from 0.0247 to 0.0302 while correlation declined to 0.587. At 50% input feature-map dropout, standard deviation rose to 0.0315 while correlation declined to 0.517 (Figure 5G). Posterior averaging did not make the point estimate more robust; the result supports uncertainty awareness rather than robustness through ensembling.

## 4. Discussion

### 4.1. A representation-centered account of rest-to-task prediction

In 40 held-out healthy participants from the WU-Minn HCP, VRPTR improved prediction of individual language-task activation from resting-state connectivity and secondary analyses indicated improved recovery of participant-specific deviations from the training-derived population template. The result extends evidence that intrinsic network architecture constrains task states and that activity flow over resting-state networks shapes task activation (Cole et al., 2014; Cole et al., 2016), together with direct rest-to-task prediction studies (Ngo et al., 2022; Tavor et al., 2016), by separating shared task anatomy from individualized structure. Whole-map evaluation alone would have obscured that distinction, since the training-derived template was already a strong predictor, yet VRPTR recovered additional residual structure and separated the correct participant from the nearest alternative more strongly than either comparator.

This focus on representation follows the original motivation for VRPTR (Di Giovanni, 2024) and the broader clinical argument that resting-state models must accommodate individual functional organization (Di Giovanni & Collins, 2023). The present results refine earlier findings; we determine that individualization emerges from the interaction between a shared transformed bottleneck and spatially resolved multiscale pathways.

### 4.2. Transformer processing contributed to prediction accuracy

The factorial experiment supports a contribution of Transformer processing in this architecture. Removing the Transformer reduced prediction and residual individualization even when spatial compression and the rest of the encoder-decoder were preserved. The deterministic Transformer model remained almost as accurate as full VRPTR and clearly outperformed both no-Transformer variants.

The architecture treats compressed channel maps as tokens, permitting global mixing among functional-connectivity features before reconstruction. Because the deepest convolutional tensor cannot bypass this stage, the ablation tests the contribution of this processing stage. Nevertheless, the swap results show that global mixing is not sufficient by itself because the tested swaps affected subject-specific activation more strongly through shallower spatial features. VRPTR works because it combines global task structure with individualized spatial conditioning.

### 4.3. The variational component should be interpreted probabilistically

Repeated latent draws produced a modest spatial association with absolute error in each test participant, supported selective prediction, and improved probabilistic scores after calibration. Uncertainty also increased under the tested input degradations. These findings support relative uncertainty information, although the mean error association was small (ρ=0.091), the probabilistic-score gains were modest, and raw intervals were severely underdispersed. Sampling conditioned on fixed weights and skip features, so it captured only one component of predictive uncertainty. The large rescaling and residual floor show that latent spread alone did not account for observed task-map error. Cross-fitting kept each participant’s target out of that participant’s calibration fit, but all folds came from the same 40-person cohort. External calibration performance, and the consequences of excluding uncertain voxels for a specific application, remain to be established.

### 4.4. Implications and limitations

Evaluation of rest-to-task models should prioritize agreement with observed activation maps. The training-derived population template supplied a fixed reference without using test outcomes. VRPTR improved spatial correlation and overlap relative to that reference, but mean MAE and RMSE were not lower. Correlation is insensitive to overall scale and offset, so improved pattern recovery does not imply improved absolute agreement. Residual maps and participant matching provide complementary evidence of specificity within this cohort; they do not establish clinical localization accuracy or uniquely different functional maps in every individual.

The exclusive focus on language is both a strength and a limitation of our study. It creates a demanding, internally coherent benchmark in which a model must recover a distributed and lateralized network rather than a single compact response, and it connects the methodological questions to applications in hemispheric-dominance assessment, eloquent-cortex localization, surgical planning, and the study of functional reorganization. These uses are especially relevant when conventional task performance is unreliable (Benjamin et al., 2017; Di Giovanni & Collins, 2023; Kumar et al., 2020). At the same time, success on language prediction does not establish task-general prediction. The architecture should next be evaluated on motor, memory, and other cognitive contrasts, and the present language-only design does not establish performance in those domains. No patients or surgical localization outcomes were evaluated; these analyses do not validate clinical language-dominance assessment.

The volumetric formulation was selected for translational compatibility. Surface representations can better respect cortical topology and reduce variability related to cortical folding (Coalson et al., 2018), whereas volumetric models provide direct compatibility with routine image volumes and retain non-cortical locations. A future comparison using matched surface and volumetric implementations would clarify the consequences of this representational choice, since the controlled comparators were volumetric adaptations rather than exact reproductions of published surface-based architectures. Atlas choice can influence the connectivity features, and registration, smoothing, and mask definition can affect voxelwise overlap even though predicted activation is not constrained to parcel boundaries. Moreover, replacing the Transformer with identity mappings removes trainable processing capacity, so these ablations do not isolate attention from parameter count or compute. The current comparison also does not establish superiority to an optimized linear or ridge-regression baseline.

Several limitations constrain generalization. The analysis used one harmonized healthy-adult dataset, one language contrast, a 40-participant evaluation cohort, and one matched training seed. Participant IDs were disjoint across training and evaluation, but family-wise separation was not established; HCP relatedness could therefore affect generalization estimates and participant-level inference. Moreover, the study used a fixed subset rather than all eligible HCP participants. This design strengthened the control of the model and ablation comparisons but limits conclusions about performance scaling with additional training data. Replication using the remaining eligible HCP participants, or preferably an independent acquisition cohort, will be needed to determine whether the observed architectural effects persist at larger sample sizes and across datasets. The observed task map was treated as the reference despite known task-fMRI measurement error and potential disease-related variation in the BOLD response (D’Esposito et al., 2003; Elliott et al., 2020). The corruption experiments were synthetic, and the latent posterior omitted weight uncertainty and uncertainty transmitted through skip pathways. External datasets, repeated task acquisitions, additional cognitive domains, and clinical cohorts will be required before translational conclusions are warranted.

## 5. Conclusion

In a healthy-adult HCP language benchmark, VRPTR improved correspondence with observed task maps relative to the evaluated volumetric comparators. Matched ablations supported a contribution of Transformer processing, while component swaps implicated multiscale skip features in subject-specific spatial patterns. Variational sampling provided a modest relative uncertainty signal that required substantial calibration. These findings distinguish architectural contributions to map prediction, subject-specific activation, and uncertainty; they do not yet establish clinical utility or externally validated uncertainty estimates.

## Data and Code Availability

The healthy-control data were obtained from the WU-Minn Human Connectome Project and are available through https://www.humanconnectome.org/study/hcp-young-adult to users who accept the HCP data-use terms.

## Ethics Statement

The Human Connectome Project obtained informed consent and institutional approval for data acquisition, as described in the source documentation. The present work is a secondary analysis of HCP data obtained under the applicable data-use terms.

## Author Contributions

Daniel A. Di Giovanni: Conceptualization, Methodology, Software, Validation, Formal analysis, Investigation, Resources, Data curation, Visualization, Project administration, Writing – original draft, Writing – review & editing. D. Louis Collins: Supervision, Writing – review & editing.

## Acknowledgments

Data were provided by the Human Connectome Project, WU-Minn Consortium (Principal Investigators: David Van Essen and Kamil Ugurbil; 1U54MH091657) funded by the 16 NIH Institutes and Centers that support the NIH Blueprint for Neuroscience Research; and by the McDonnell Center for Systems Neuroscience at Washington University.

## Declaration of AI Assistance

ChatGPT (OpenAI) was used to assist with manuscript drafting and revision, modifications to analysis and figure-generation code, and checks of consistency between analysis outputs and reported results. The authors take full responsibility for the final manuscript, analyses, and conclusions.

